# Antiviral effect of betacyanins from red pitahaya (*Hylocereus polyrhizus*) against influenza A virus

**DOI:** 10.1101/2024.02.09.579603

**Authors:** Chie Min Lim, Sunil Kumar Lal, Nurulfiza Mat Isa, Abdul Rahman Omar, Wee Sim Choo

## Abstract

Seasonal influenza affects millions of lives worldwide, with influenza A virus (IAV) responsible for pandemics and annual epidemics, causing the most severe illnesses resulting in patient hospitalizations or death. With IAV threatening the next global influenza pandemic, it is a race against time to search for antiviral drugs. Betacyanins are unique nitrogen-containing and water-soluble reddish-violet pigments that have been reported to possess antiviral properties against dengue virus. The objective of this study was to examine the antiviral effect of betacyanins from red pitahaya (*Hylocereus polyrhizus*) on IAV-infected lung epithelial A549 cells. HPLC and LC-MS analysis of extracted betacyanin showed four betacyanins in the betacyanin fraction, namely phyllocactin, hylocerenin, betanin, and isobetanin. Cytotoxicity assay showed that betacyanin fractions were not cytotoxic to A549 cells at concentrations below 100 µg/mL. Betacyanin fraction concentrations of 12.5, 25.0, and 50.0 µg/mL prevented the formation of viral cytopathic effect and reduced virus titer in IAV-infected cells up to 72 h. A downregulation of protein and mRNA nucleoprotein expression levels was observed after treatment with 25.0 and 50.0 µg/mL of betacyanin fraction after 24 h, thereby providing evidence for the anti-viral activity of betacyanin from red pitahaya against IAV *in vitro*.

## Introduction

The influenza virus is an enveloped RNA virus in the Orthomyxoviridae family of which; the virus breaks down into four classes of the Orthomyxoviridae family - types A, B, C, and D, with influenza A virus (IAV) causing the most severe illnesses while being responsible for its annual epidemics and pandemics resulting in patient hospitalizations or death [1]. In total, eight ribonucleic acid (RNA) segments build the IAV genome, of which they code for hemagglutinin (HA), M1, M2, neuraminidase (NA), nucleoprotein (NP), nonstructural protein (NS) 1, NS2, polymerase acidic protein (PA), polymerase basic (PB)1, PB1-F2, and PB2 [2]. The World Health Organization (WHO) reported approximately 1 billion cases of seasonal influenza annually, of which 3 to 5 million cases are severe and can cause up to 650,000 deaths annually [3]. Additionally, global influenza pandemic is being listed as one of the ten threats to global health in 2019 [4]. Moreover, the WHO also stated that a worldwide influenza pandemic is on the horizon, and the only uncertain thing is when it will hit and how severe it will be [4]. Currently, the National Institute of Health (NIH) lists oseltamivir (Tamiflu), peramivir (Rapivab), and zanamivir (Relenza) as recommended antiviral drugs to treat IAV infection [5]. While these drugs are currently adequate, due to the nature of the drugs targeting a key viral factor NA, the ability of the virus to develop resistance to antiviral therapies is inevitable. Resistant strains reduced the susceptibility to oseltamivir [6] and two other drugs, rimantadine (Flumadine) and amantadine (Symmetrel) that are used to treat IAV infection were no longer recommended in the United States [5]. More worryingly, resistant strains have been detected in immunocompromised patients [7].

Betacyanin is a reddish-violet pigment that belongs to a class of plant pigments called betalains [8]. The Cactaceae families (namely, *Hylocereus* sp. and *Opuntia* sp.) and the Amaranthaceae families (namely, *Amaranthus* sp. and *Beta vulgaris* L.) are two of the most common and well-known sources of betacyanin [8]. Betalains have been well-documented to exhibit antibacterial activity against a broad range of microbes, most notably against multidrug-resistant bacteria and biofilm-producing bacteria [9]. Numerous *in vivo* and *in vitro* studies have suggested the numerous biological benefits of betacyanins from a wide range of fruits and vegetables, such as red pitahaya or red dragon fruit (*Hylocereus polyrhizus*), prickly pear (*Opuntia ficus-indica*), and red beetroot (*B. vulgaris*). Some of these benefits include anti-inflammatory responses [10, 11], protection against DNA damage [12, 13], and stabilizing cell cycle regulations [14]. Despite the broad antibacterial properties of betacyanin, as well as the numerous health benefits from the mechanism studies, none of these studies were performed on viral models. Only one study demonstrated the antiviral effect of betacyanin from red pitahaya and red spinach against the dengue virus serotype 2, showing that betacyanin was able to reduce virus titers in the form of plaque reduction [15]. This study aimed to investigate the antiviral effect of betacyanins from red pitahaya (*Hylocereus polyrhizus*) on IAV-infected lung epithelial A549 cells.

## Methods

### Betacyanin extraction, semi-purification & quantification

Red pitahaya fruits (*H. polyrhizus*) were purchased from a local supermarket (Aeon, Malaysia). To prepare for betacyanin extraction, the skin was first peeled, and the flesh was then blended with ice-cold 70% methanol in a ratio of 4 L of 70% methanol for every 1 kg of fruits used with a commercial blender. The blended pulp was centrifuged at 2,000 × g, 4°C for 10 min, and the supernatant was filtered under vacuum. The crude betacyanin extract was concentrated using a rotary evaporator, freeze-dried at −80°C, and kept at −80°C until the next step.

Semi-purification of the crude betacyanin extract was carried out as described by another study via column chromatography using Amberlite XAD16N polyaromatic adsorbent resin (Sigma Aldrich, USA) [15]. Briefly, 1 L of distilled water was first used to wash the packed column containing the adsorbent resin. Next, 500 mL of 2% sodium hydroxide solution was used to activate the resin. After activation, distilled water was used to wash the sodium hydroxide, and acidified water prepared with trifluoroacetic acid (Sigma Aldrich, USA) was used to condition the pH to 3.0. Once conditioned, 10 mL of Milli-Q water was used to dissolve 5 g of dried crude betacyanin extract, added to the column, and then desalted with 0.5 L of the same acidified water as mentioned previously. Next, elution with 0.2 L of acidified methanol prepared in a 95:5 ratio of methanol to acidified water was carried out. The eluted fraction was collected and concentrated using a rotary evaporator, freeze-dried at −80°C, and kept at −80°C until further use. This dried fraction was termed the betacyanin fraction.

Betacyanin content was quantified according to a method published by another study with slight modifications [16]. Absorbance reading was obtained via a spectrophotometric method using a Lambda 25 UV-Vis spectrophotometer (Perkin Elmer, USA) at a wavelength of 538 nm. The betacyanin content in the betacyanin fraction was calculated as follows:

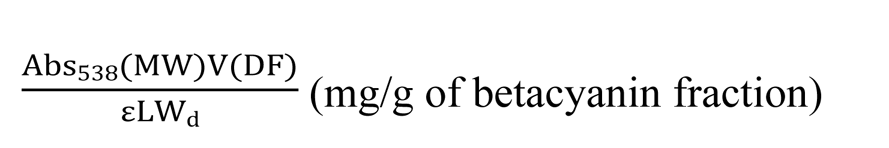

Where Abs_538_ = absorbance at 538 nm; MW = 550.473 g/mL; V = volume of betacyanin fraction solution (mL); DF = dilution factor; ε = 65,000 cm^-1^ mol^-1^ L; L (path length) = 1.0 cm; W_d_ = weight of betacyanin fraction (g) Before using the reconstituted betacyanin in any experiments, the reconstituted betacyanin was filter-sterilized through a 0.22 µM pore-sized membrane.

### Betacyanin characterization

A high-performance liquid chromatography (HPLC) analysis was performed according to another study [17], involving the use of an Agilent 1260 Infinity HPLC system (Agilent Technologies, USA) equipped with a Purospher® STAR RP-18 end-capped LiChroCART® column (Merck, Germany) and a photodiode array detector. Briefly, 8 mg/mL of betacyanin fraction was prepared using Milli-Q water, filtered through a 0.45 μM pore-sized membrane, and 20 μL of the sample was injected into the column. The mobile phase cocomprised00 mL of acetonitrile (Sigma Aldrich, USA) and 900 mL of 0.5% trifluoroacetic acid (Sigma Aldrich, USA). A 0.5 mL/min flow rate under isocratic mode was applied for approximately 50 min with detection set at 538 nm. The percentage of betacyanin was calculated by dividing the individual peak area by the total peak area. A liquid chromatography-tandem mass spectrometry (LC-MS/MS) analysis was performed to characterize the betacyanin fraction further using an Agilent 6520 Accurate Mass Q-TOF LC/MS (Agilent Technologies, USA) operating in a positive ionization mode with dual electrospray ionization source [18].

### Cell culture preparations

Human lung adherent epithelial A549 cells (CCL-185, ATCC, USA) and cocker spaniel kidney adherent epithelial MDCK cells (NBL-2, CCL-34, ATCC, USA) were both routinely cultured in high-glucose Dulbecco’s Modified Eagle Medium (DMEM, Gibco, USA) supplemented with 10% fetal bovine serum (Tico Europe, Netherlands) without antibiotic use. A549 cells were used for all infection experiments, while MDCK cells were used for plaque assay only as MDCK cells have a high affinity for influenza virus and form distinct plaques [19]. Cells were subcultured as needed. Both cell lines were grown in a 37°C humidified incubator supplied with 5% CO_2_. *Mycoplasma* contamination was screened routinely via a polymerase chain reaction (PCR)-based approach.

### A549 host cell cytotoxicity evaluation

The cytotoxicity of betacyanins and oseltamivir phosphate (FluHalt, India) on A549 cells was evaluated using 3-(4,5-dimethylthiazol-2-yl) −2,5-diphenyltetrazolium bromide (MTT) assay. Oseltamivir phosphate was to be used as a positive control for all subsequent downstream assays that involved influenza infection and as such, a suitable concentration was needed to be determined via the MTT assay approach. A549 cells were first seeded in a 96-well plate for 24 h prior to the assay. A range of betacyanin fractions (1.56, 3.13, 6.25, 12.5, 25, 50, and 100 µg/mL) or oseltamivir phosphate (50, 100, 200, and 400 µM) was added to the cells, and incubated for another 24 h in a humidified incubator set to 37°C with 5% CO_2_. Additionally, a positive control of cytotoxicity for this specific assay was also included by adding 1% Triton X-100 (Amresco, USA) to facilitate the lysis of cells. Once incubated, the A549 cells were washed once with 1× phosphate-buffered saline (PBS) before incubating again with 100 µL of 0.5 mg/mL of MTT reagent (Merck, Germany) for 3 h at 37°C to allow the formation of purple formazan. To facilitate the solubilization of the formazan precipitates, 200 µL of dimethyl sulfoxide (DMSO) (Vivantis, Malaysia) was added and incubated at 37°C for an additional 30 min. Absorbance reading was obtained via a spectrophotometric method using an Infinite M Nano+ microplate reader (Tecan, Switzerland) at a wavelength of 630 nm for the background and 570 nm for the signal. A graph of cell viability against betacyanin fraction was plotted using GraphPad Prism (Version 9.3, GraphPad, USA) to identify the suitable betacyanin fraction and oseltamivir phosphate concentration for downstream assays.

### Virus infection

A549 cells to be infected with influenza A virus (VR-95, ATCC, USA) were firstly seeded and incubated in a 12-well plate for 16 h and then infected at a multiplicity of infection (MOI) of 1 for 1 h in the presence of infection media (DMEM supplemented with 0.3% bovine serum albumin fraction V) with gentle rocking every 15 min in 37°C at 5% CO_2_. The virus inoculum was discarded after 1 h, and the infected cells were washed once with 1× PBS. A virus growth medium consisting of infection media supplemented with an additional 1 µg/mL trypsin-TPCK (ThermoFisher, USA) was added to the cells. In addition to the virus growth medium, three groups of cells received an additional 12.5, 25.0, and 50.0 µg/mL of betacyanin fraction, the positive control received an additional 100 µM of oseltamivir phosphate, and the negative control group received only virus growth medium. All groups were incubated for 24 h at 37°C. The 24 h time point was chosen as it was shown in published literature from our laboratory that 24 h was sufficient to induce cellular responses due to virus infection [20, 21]. After incubation, the supernatant was harvested, aliquoted, and stored at −80°C until use for plaque assay. TRIzol reagent (Invitrogen, USA) was added to the cells for RNA and protein isolation.

### Plaque assay

MDCK cells were first seeded in 6-well plates for 16 h. Cells were then infected with supernatants diluted at various dilution factors with infection media with gentle rocking every 15 min for 1 h at 37°C at 5% CO_2_. The virus inoculum was discarded after the 1 h incubation period, and cells were washed once with 1× PBS. The cells were overlayed with a mixture containing 0.3% low-melting point agarose (ThermoFisher, USA), L-15 media (HiMedia, India), 20 mM HEPES (Invitrogen, USA), 0.075% sodium bicarbonate (Merck, USA), and 2 µg/mL trypsin-TPCK. Cells were then incubated at 37°C at 5% CO_2_ for 48 h. The overlay media was discarded, and a crystal violet solution (0.1% crystal violet (Merck, Germany), 0.04% ethanol, 20% methanol) was used to fix and stain the cells for 30 min in the dark at room temperature. After discarding the crystal violet solution, the cells were rinsed briefly with distilled water. The stained cells were allowed to dry, and the number of plaques visible under the naked eye was counted. The virus titer was calculated with the following formula expressed as plaque forming unit/milliliter (pfu/mL): Virus titer (pfu/mL) = Number of plaques / [Dilution factor × volume added (mL)]

### Viral CPE analysis

The direct antiviral effect of betacyanin fractions was evaluated through the formation of viral CPE using the Viral ToxGlo^TM^ Assay (Promega, USA) per the manufacturer’s instructions with slight modifications. A549 cells were first seeded in a 96-well plate for 16 h. Infection and betacyanin and positive control treatments were performed, as mentioned previously. Infection media replaces virus inoculum in mock-infected cells. Prior to the addition of the ATP Detection Reagent, an inverted phase contrast microscope (CKX41, Olympus, Japan) with an attached digital camera (DP21, Olympus, Japan), connection kit (DP21-SAL, Olympus, Japan), and hand switch (DP21-HS, Olympus, Japan) was used to capture microscopic images of the cells to visualize any morphological changes. The treatment media were aspirated and discarded after 24 h. PBS and ATP Detection Reagent were mixed in a 1:1 ratio and added to the cells. The cells were incubated at room temperature for 30 min on a shaker at 35 rpm. Relative light unit (RLU) was obtained via a luminescence method using an Infinite M200 microplate reader (Tecan, Switzerland). A graph of RLU against betacyanin fractions was plotted using GraphPad Prism (Version 9.3, GraphPad, USA).

### RNA and protein isolation

Total RNA isolation was performed as stated on the manufacturer’s instructions using the TRIzol reagent. Briefly, TRIzol reagent was added to the cells and resuspended. Chloroform (Merck, Germany) was added to the mixture, inverted, and incubated for 5 min before centrifuging at 12,000 × g, 4°C for 15 min. The top aqueous layer was transferred to a new centrifuge tube, leaving behind the middle interphase and lower organic layers. Isopropanol (Merck, Germany) was added to the centrifuge tube containing the aqueous layer and was incubated for 10 min to precipitate the RNA. The lower organic layer was kept for protein isolation in the next step. The mixture was centrifuged at 12,000 × g, 4°C for 10 min. The supernatant was discarded, the RNA pellet was washed in 75% ethanol (Systerm, Malaysia), and then centrifuged at 7,500 × g, 4°C for 5 min. The supernatant was discarded, and the pellet was air-dried for 10 min. The pellet was resuspended in RNA solubilization buffer (20 mM dithiothreitol (Vivantis, Malaysia) and 1 mM sodium citrate (Merck, Germany). The yield and purity were determined with a BioDrop Spectrophotometer Duo (Biochrom, UK), and the integrity was determined via agarose gel electrophoresis. The RNA was stored at −80°C until needed.

To the lower organic layer, 100% ethanol was added, incubated for 5 min, and centrifuged at 2,000 × g, 4°C for 5 min. The protein was precipitated from the supernatant by adding isopropanol for 10 min after transferring the supernatant to a new tube. The mixture was centrifuged at 12,000 × g, 4°C for 10 min. The protein pellet was washed in protein wash buffer (0.3 M guanidine hydrochloride (Sigma Aldrich, USA) in 95% ethanol) for 20 min after discarding the supernatant, followed by centrifugation at 7,500 × g, 4°C for 5 min. The wash buffer was discarded, and the washing step was repeated twice. A final wash was performed by incubating in 100% ethanol for 20 min and centrifugation at 7,500 × g, 4°C for 5 min. The pellet was air-dried for 10 min after discarding the final ethanol wash. The pellet was resuspended in protein resuspension buffer (150 mM NaCl (Merck, USA), 5 mM EDTA (Merck, USA), 10 mM Tris-HCl pH 8.0 (Merck, USA), and 1% SDS (Nacalai Tesque, Japan). The pellet was incubated at 55°C for 10 min to facilitate the pellet dissolving into the buffer. The mixture was centrifuged at 10,000 × g, 4°C for 10 min. The supernatant was transferred to a new tube, and protein yield was quantified using bicinchoninic acid (BCA) assay. The quantified protein was kept at −80°C until needed.

### Viral NP mRNA expression analysis

Reverse transcription of the total RNA isolated to complimentary DNA (cDNA) was carried out using the ReverTra Ace qPCR RT Master Mix with gDNA Remover kit (Toyobo, Japan) per the manufacturer’s instructions. Briefly, 200 ng of total RNA was denatured at 65°C for 5 min. Next, 4× DN Master Mix was added to remove genomic DNA contaminants and incubated at 37°C for 5 min. Lastly, reverse transcription was performed by adding 5× RT Master Mix II and then incubating at 37°C for 15 min, followed by 50°C for 5 min, and 98°C for 5 min. The synthesized cDNA was stored at −20°C until needed.

Real time PCR (qPCR) was performed with the PrecisionPLUS qPCR Master Mix (Primerdesign, UK) per the manufacturer’s instructions. Briefly, 2 µL of synthesized cDNA was firstly diluted in a 1:5 ratio and used in a 20 µL reaction volume using the CFX96 Touch Real-Time PCR Detection System (BioRad, USA) with the following thermal cycling conditions: 95°C for 2 min, followed by 40 cycles of 95°C for 10 s, 60°C for 60 s. Data were collected at the end of each cycle. A melt curve analysis was also performed at the end of the thermal cycling condition to verify product formation. A non-template control (NTC) was included to verify the sterility of the reagents used. Data were analyzed using the CFX Maestro Software (BioRad, USA). The primer sequences are listed below:

<u>Influenza A virus mRNA NP [22]</u>

Forward: 5′ – CTCGTCGCTTATGACAAAGAAG – 3′

Reverse: 5′ – *AGATCATCATGTGAGTCAGAC* – 3′

<u>β-actin</u>

Forward: 5’ – CTTCGCGGGCGACGAT – 3’

Reverse: 5’ – CCACATAGGAATCCTTCTGACC – 3’

### Viral NP protein expression analysis

The proteins isolated previously were first boiled at 100°C for 5 min in the presence of a reducing agent, 2-mercaptoethanol (Sigma Aldrich, USA). Next, 50 µg of the boiled and reduced protein was subjected to sodium dodecyl sulfate–polyacrylamide gel electrophoresis (SDS-PAGE) on a 12.5% resolving gel and 5% stacking gel and subsequently transferred to nitrocellulose membranes (Amersham, United Kingdom). The membranes were incubated with 5% skim milk (Nacalai Tesque) in PBST (0.1% Tween-20 (Vivantis, Malaysia) in PBS) on a rocker at 35 rpm for 60 min at room temperature to prevent nonspecific binding. The blots were then probed with influenza A virus nucleoprotein antibody (GTX125989, GeneTex, USA) (1:1500 dilution in 5% skim milk) at 4°C overnight on a rocker at 35 rpm. The blots were washed three times with PBST and then incubated for 120 min at room temperature on a rocker at 35 rpm with the corresponding horseradish peroxidase-conjugated secondary antibody (1:5000 dilution in 5% skim milk). The blots were washed thrice with PBST, and the signals from the reactions were detected using enhanced chemiluminescent (Merck, Germany) and visualized using G:Box Chemi XX6 Gel Viewer (Syngene, United Kingdom). As an internal control, vinculin was used (sc-73614, Santa Cruz Biotechnology, USA) (1:1000 dilution in 5% skim milk), and all protein quantities were normalized to the intensities of the vinculin bands. NP and vinculin expression levels were subjected to densitometric analysis using ImageJ (Version 1.8.0, National Institutes of Health, USA).

### Time-point virus infection & viral CPE formation analysis

A549 cells were first seeded and incubated in a 12-well plate for 16 h. Infection and betacyanin and positive control treatments were performed, as mentioned previously, with slight modifications. Here, cells were only given 50.0 µg/mL of betacyanin fraction, as previous data suggested 50 µg/mL of betacyanin fraction showed the highest level of antiviral activity. All groups were incubated for 48 or 72 h at 37°C. After incubation, the supernatant was subjected to plaque assay.

Similarly, a time-point study was also conducted on the formation of viral CPE. A549 cells were first seeded and incubated in a 96-well plate for 16 h. Virus infection and 50 µg/mL of betacyanin and positive control treatments were performed as mentioned previously. Microscopic images were taken, and luminescence reading was acquired, as mentioned previously.

### Data analysis

All experiments were performed in biological triplicates. The fold-change in NP mRNA expression between groups was analyzed using the delta-delta Ct method. All data were expressed in mean value ± standard deviation. Data collected from performed assays were analyzed using one-way ANOVA and Tukey’s post-hoc test using GraphPad Prism to determine statistical significance between groups. A p-value lesser than 0.05 was considered significant.

## Results and discussion

### Characterization of betacyanins

Table 1 shows the characterization of betacyanins in the betacyanin fraction extracted from red pitahaya. The known betacyanins identified in the betacyanin fraction of red pitahaya were betanin, isobetanin, phyllocactin, and hylocerenin (Table 1, and supplementary information, Fig S1 – S7). The most abundant betacyanin was phyllocactin (57.07 ± 0.25%), followed by betanin (22.21 ± 0.11%), hylocerenin (12.18 ± 0.10%), and finally, isobetanin (4.73 ± 0.10%) (Table 1). This composition is in accordance with previous studies [15, 18, 23].

**Table 1.**
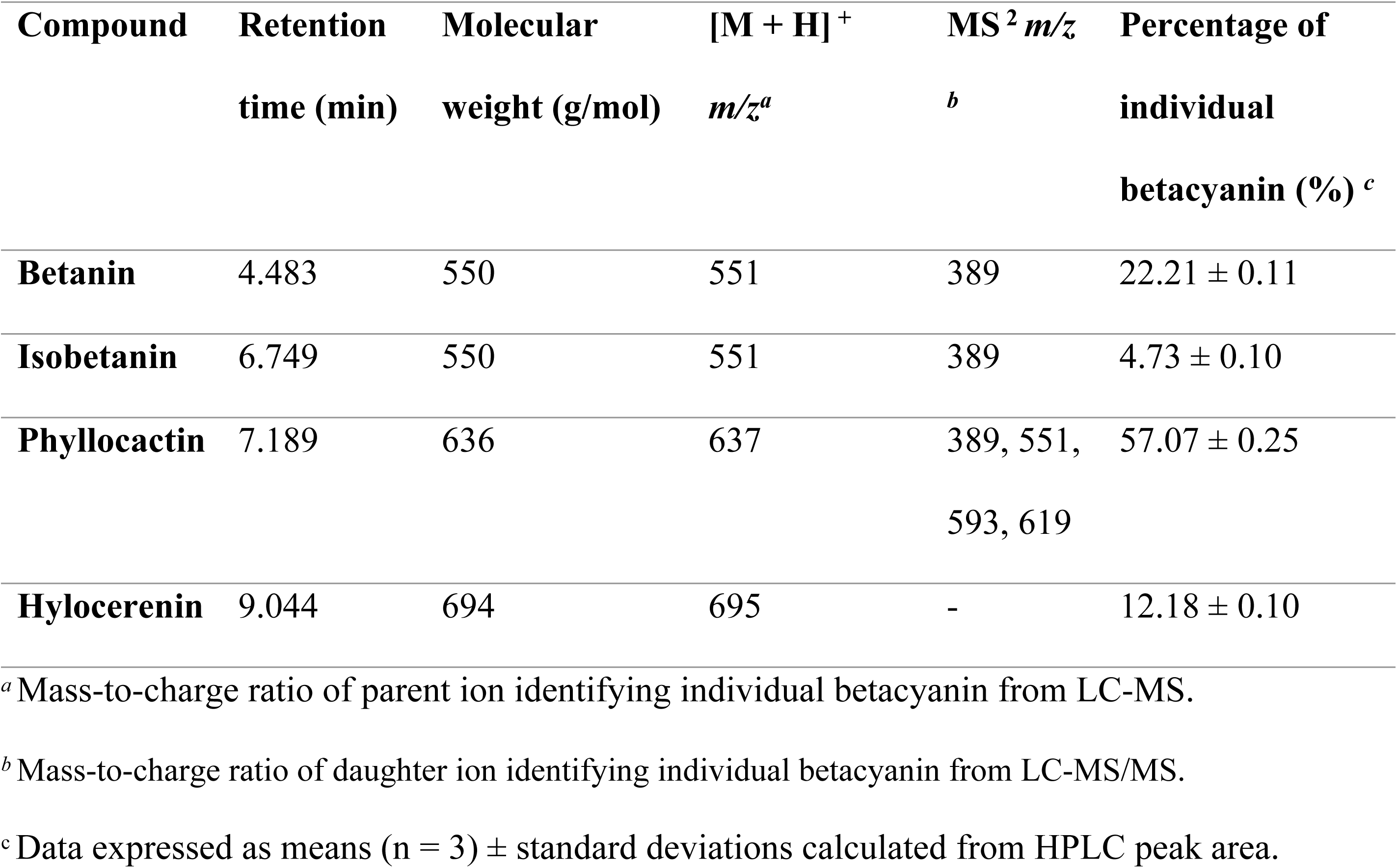
HPLC and LC-MS/MS profile of betacyanins from red pitahaya.

### Betacyanins and oseltamivir phosphate were not cytotoxic to A549 cells

The cell viability of A549 cells was reduced by 19.3%, 2.0%, 11.8%, 16.5%, 19.6%, 31%, and 89.7% after being treated with 1.56, 3.13, 6.25, 12.5, 25, 50 and 100 µg/mL of betacyanin fraction, respectively (Fig 1). In pursuant to ISO 10993-5:2009 published by the International Organization for Standardization (ISO) that describes the methodology to determine the range of which a compound is considered cytotoxic *in vitro* [24], all selected betacyanin concentrations, except for 100 µg/mL, were not cytotoxic to A549 cells, even though the cell viability in A549 cells treated with 50 µg/mL (p-value = 0.0322) was significantly reduced. The standard published by ISO indicated that not more than 50% of cell death is considered non-cytotoxic [24], with several published literature utilizing this metric as well [25, 26]. Thus, concentrations of 12.5, 25, and 50 µg/mL of betacyanin fraction were chosen for downstream experiments. This is the first study that examined the cytotoxicity of betacyanins from red pitahaya on A549 cells. Another study has reported that the half maximum cytotoxicity concentration of betacyanins from red pitahaya was 664.73 µg/mL in monkey kidney epithelial Vero cells [15], while another study reported no observed cytotoxicity exerted on both human embryonic kidney HEK293 cells and THP-1 monocytes at a concentration as high as 6.25 mg/mL [17]. The cell viability of A549 cells was reduced by 6.4%, 12.7%, 9.0%, and 4.9% after being treated with 50, 100, 200, and 400 µM of oseltamivir phosphate, respectively. As described above, none of the selected oseltamivir phosphate concentrations were cytotoxic to A549 cells in accordance with ISO 10993-5:2009.

**Fig 1.**
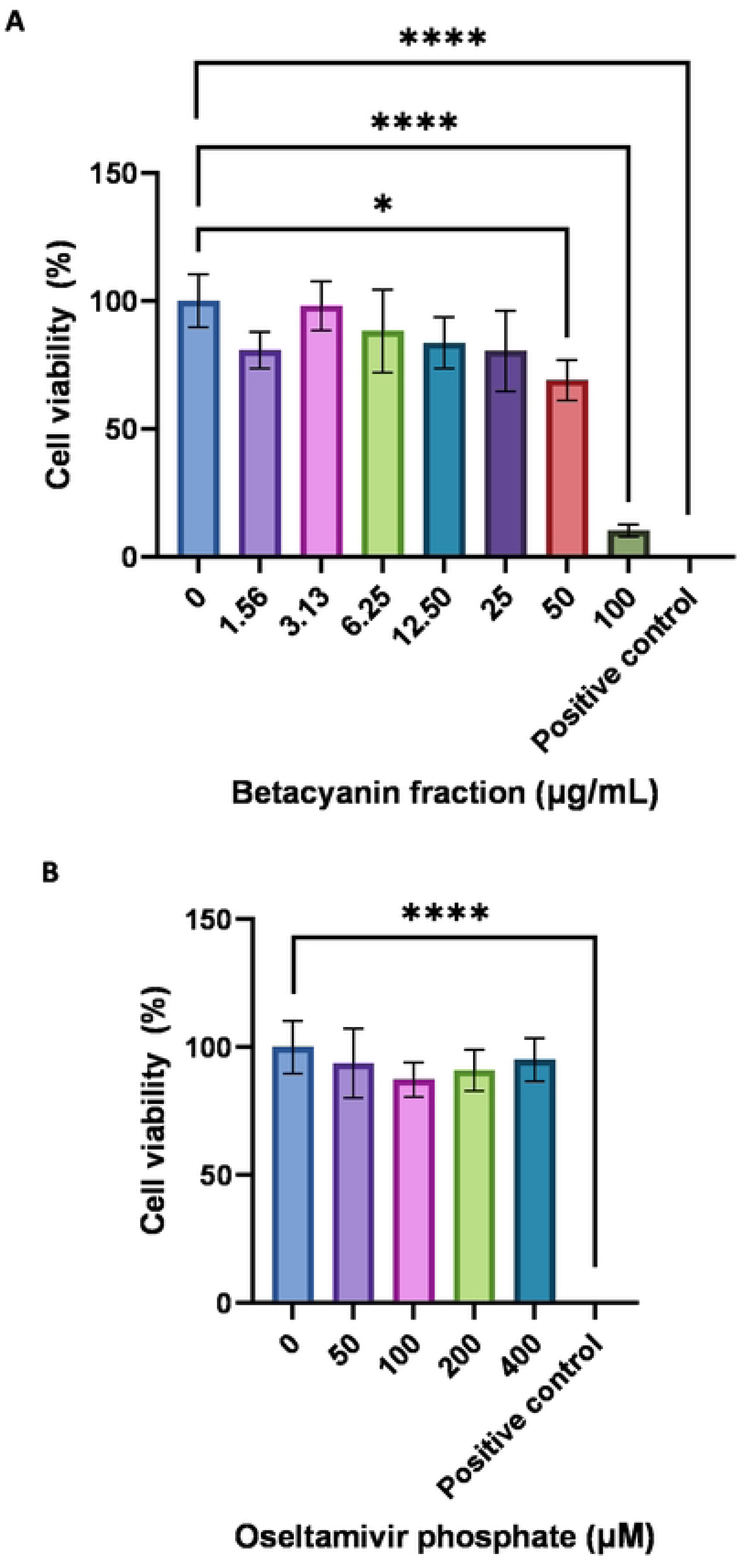
Cell viability of A549 cells at various betacyanin fraction (A) and oseltamivir phosphate (B) concentrations. A549 cells were first seeded in a 96-well plate. The cells then received various betacyanin fraction or oseltamivir phosphate concentrations diluted with complete media for 24 h. The positive control consisted of 1% Triton X-100 in complete media. MTT reagent was added to the cells after 24 h, and then absorbance value was acquired using a spectrophotometer. Data were presented as means (n =3) ± standard deviations and were subjected to an independent sample t-test to determine if changes in cell viability between untreated and treated ranges were significant. Statistical significance was denoted by an asterisk(*), where* has a p-value < 0.05, and**** has a p-value < 0.0001.

### Betacyanins prevented the formation of viral CPE in IAV-infected A549 cells

Infection by cytocidal viruses like IAV is usually associated with many changes in cell physiology and biochemistry, of which the formation of CPE is a sign of morphological change due to IAV infection [27]. Measurement of ATP depletion in the form of relative light unit (RLU) can be performed and correlated with viral CPE formation. Here, we are interested in examining if betacyanins can prevent the formation of viral CPE, which translates to antiviral responses.

When compared to the untreated group without any treatment, 12.5, 25.0, and 50.0 µg/mL of betacyanin content increased RLU by 22.2%, 48.2%, and 59.9%, respectively (Fig 2). The increase in RLU values indicated that betacyanin prevented the formation of viral CPE induced by IAV in A549 cells. The tested betacyanin contents exhibited direct antiviral effects, of which 25.0 (p-value = 0.0016) and 50.0 µg/mL (p-value = 0.0003) showed statistical significance. Interestingly, compared to the mock-infected group, the increase in RLU in 12.5 (p-value = 0.0223), 25.0 (p-value = 0.0002) and 50.0 µg/mL (p-value = <0.0001) betacyanin fraction groups was significant, suggesting that betacyanin may possess protective effects against trypsin that is present in the virus growth medium. Similarly, compared to the positive control, both 25.0 (p-value = 0.0014) and 50.0 µg/mL (p-value = 0.0002) betacyanin fraction groups possessed a significantly higher RLU value, suggesting that betacyanin may have further interactions with other host factors in the infected cells that may be advantageous when compared to the positive control, as shown by existing literature.

**Fig 2.**
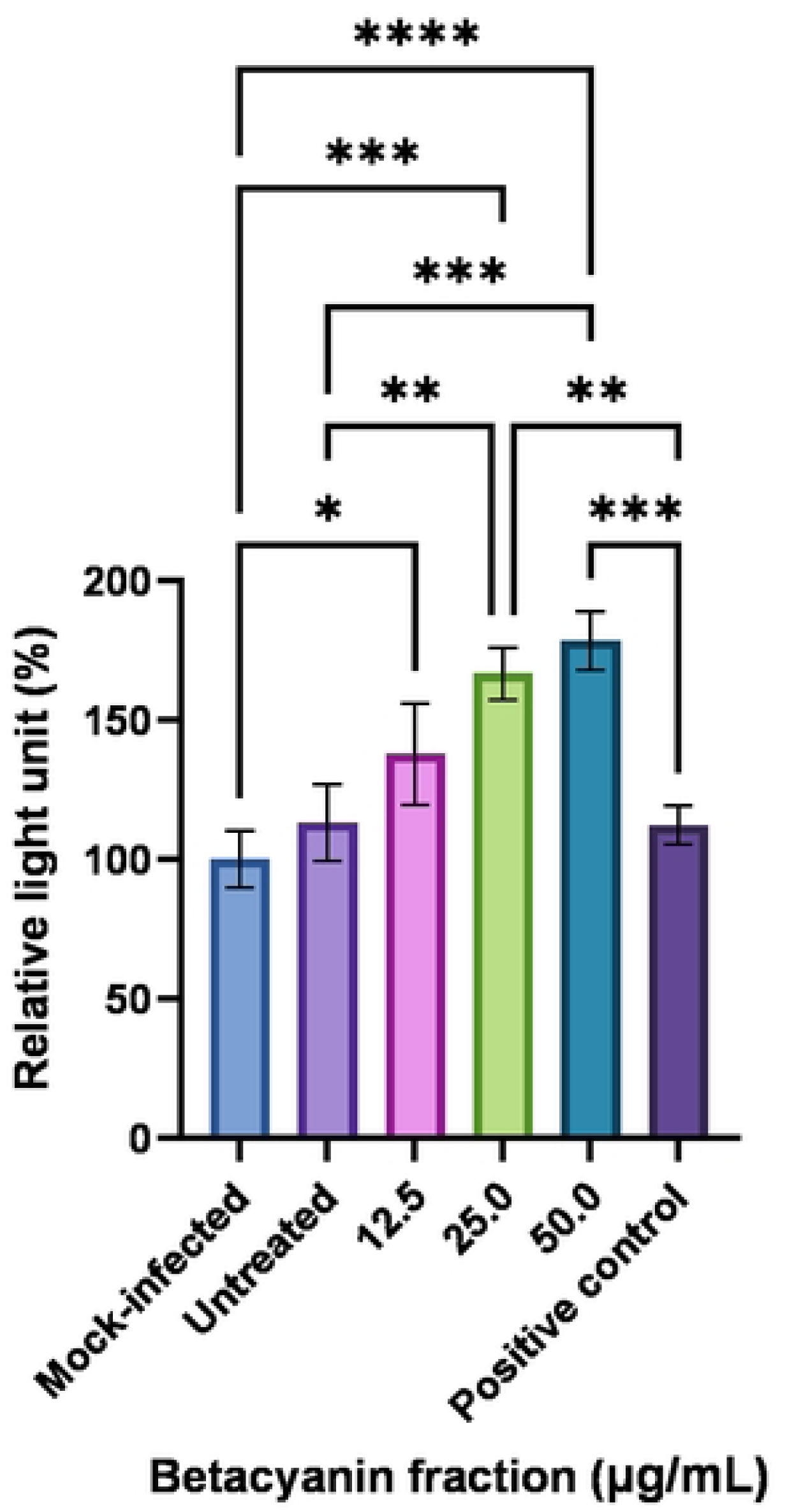
Relative light unit (RLU) after normalizing to the untreated group in IAV-infected A549 cells after treatment with various betacyanin fraction concentrations. A549 cells were first seeded in a 96-well plate and then infected at an MOI of I with IAV. The cells received various concentrations of betacyanin fraction after infection for 24 h. The positive control group received I00 µM of oseltamivir phosphate. All groups received virus infection, while the mock­ infected group received virus growth medium only without virus. After 24 h, the treatment media were removed and replaced with ATP Detection Reagent. RLU was acquired via a luminescence method. Data were presented as means (n = 3) ± standard deviations and were subjected to one­ way ANOVA to determine if changes in RLU between groups were significant. Statistical significance was denoted by an asterisk(*), where* has a p-value < 0.05, ** has a p-value < 0.01, *** has a p-value of<0.001, and**** has a p-value of<0.0001.

Morphologically, A549 cells infected with IAV demonstrated near-complete destruction, losing their distinct cellular shape (Supplementary information, Fig S8c) compared to the cells before any infection and treatment (Fig S8a). Similarly, mock-infected cells lost their distinct cellular shape (Supplementary information, Fig S8b) despite not receiving any virus treatment. Infected cells that received 12.5, 25.0, and 50.0 µg/mL of betacyanin fraction retained their morphological shape, indicating viral CPE formation was prevented (Supplementary information, Fig S8e – g). Comparing the positive control group and the betacyanin-treated groups, the betacyanin-treated groups retained their morphological shape better than the positive control, as partial destruction of cells can still be seen in the positive control group (Supplementary information, Fig S8d). This is reflective of the RLU values, as discussed previously (Fig 2). Here, betacyanin demonstrated a strong antiviral response against IAV-infected cells by preventing the formation of viral CPE and retaining the original cellular morphology.

### Betacyanins reduced virus titers in IAV-infected A549 cells

At betacyanin fraction concentrations of 12.5, 25.0, and 50.0 µg/mL, virus titers were found to be at 10800, 10933, and 9333 pfu/mL, respectively (Fig 3a), or a reduction of 23.6%, 22.6% and 34.0% in virus titer, respectively (Fig 3b). All betacyanin fractions exhibited antiviral activity by directly reducing the virus titers in IAV-infected cells. However, only the reduction in virus titer treated with 50 µg/mL of betacyanin fraction was found to be statistically significant (p-value = 0.0464). On the other hand, the positive control’s virus titer was 1173 pfu/mL (Fig 3a), or a reduction of 91.8% in virus titer (Fig 3b) (p-value = <0.0001). Oseltamivir phosphate is a commonly used antiviral agent against IAV infections as it inhibits the action of NA, thus preventing the release of viral progenies from infected cells. Mock-infected cells exhibited no visible plaques (Supplementary information, Fig S9a). A decrease in a number of visible plaques can be observed when infected cells were treated with 12.5, 25, and 50 µg/mL of betacyanin, as well as the positive control group (Supplementary information, Fig S9b - f).

**Fig 3.**
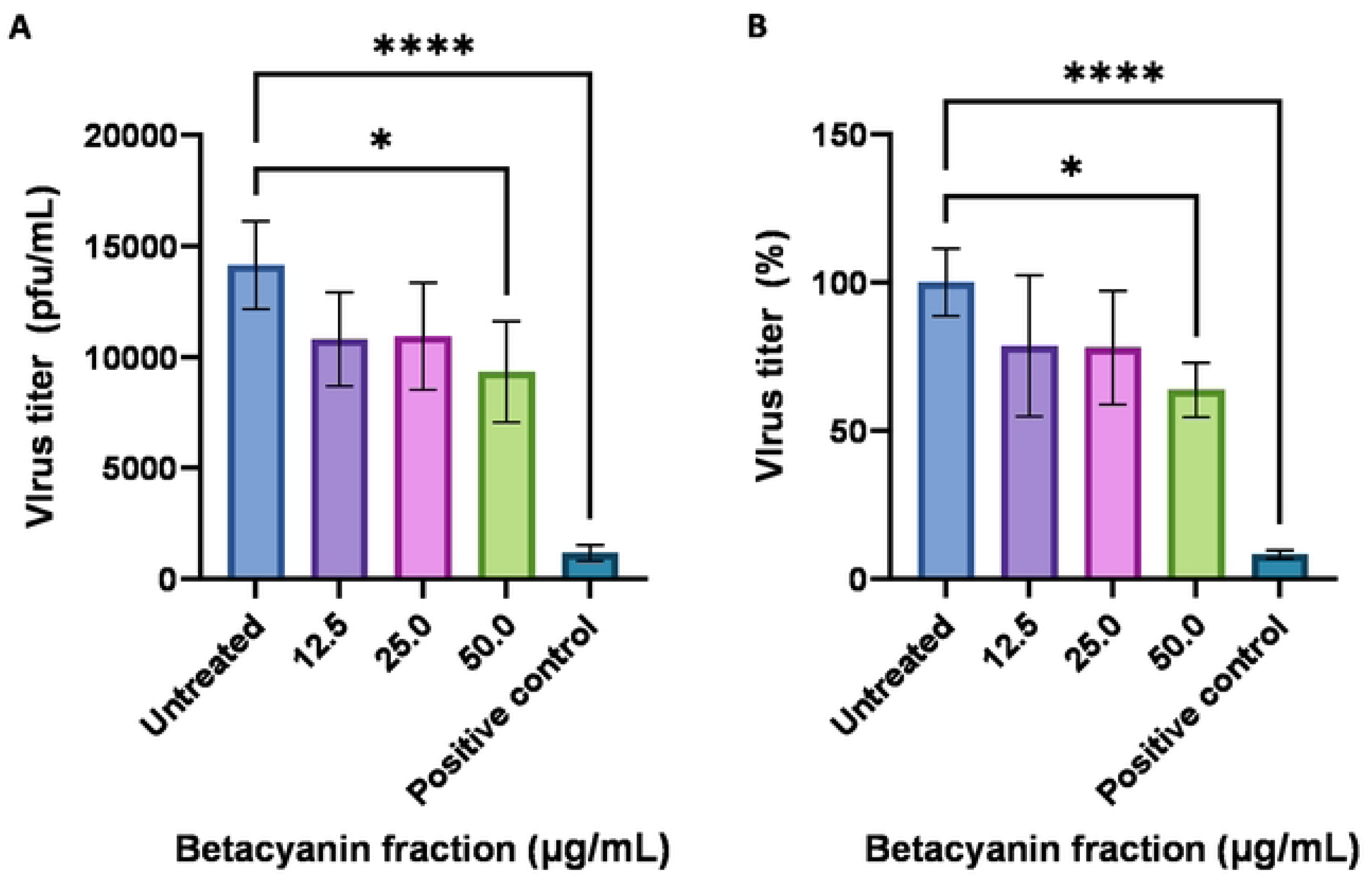
Virus titer expressed in pfu/mL (A) and percentages (B) in IAV-infected A549 ceUs after treatment with various betacyanin fraction concentrations. A549 cells were first seeded in a 12-well plate and then infected at an MOI of I with IAV. The cells received various concentrations of betacyanin after infection for 24 h. The positive control group received 100 µM of oseltamivir phosphate. The supernatant was harvested after 24 h and was subjected to plaque assay. MDCK cells were seeded in a 6-well plate and then infected with the harvested supernatant at various dilution factors. The cells were then overlaid with an agarose-L-15 medium for 48 h. The agarose overlay was discarded, and cells were stained and fixed with a crystal violet solution. Visible plaques were counted, and virus titer was calculated. Data were presented as means (n = 3) ± standard deviations and were subjected to an independent sample t-test to determine if changes in virus titers between untreated and treated ranges were significant. Statistical significance was denoted by an asterisk(*), where• has a p-value < 0.05, and**** has a p-value < 0.0001.

The antiviral capability of betacyanins has been reported by another study performed on dengue virus [15]. Chang et al. [15] reported a significant reduction in virus titer of dengue virus after treatment with betacyanins extracted from red pitahaya and red spinach, with an IC_50_ value of 125.8 µg/mL betacyanins from red pitahaya. In this present study, betacyanin concentration that was higher than 50 µg/mL was not used as betacyanin will exhibit significant cytotoxicity to A549 cells and as such, an IC_50_ was not achievable. Nonetheless, this is the first study that showed that betacyanins from red pitahaya reduced the virus titer in IAV-infected mammalian cells and the second study that suggests the antiviral effect of betacyanins. Reducing virus load is an important parameter to measure as reducing virus load directly translates towards reducing viral load and disease severity in infected patients.

### Betacyanins downregulated both protein and mRNA NP expression levels in IAV-infected A549 cells

Compared to the untreated group, cells treated with 12.5 µg/mL of betacyanin fraction exhibited an upregulation of NP of 1.777 and 1.776-fold at both the mRNA and protein levels, respectively (Fig 4). On the other hand, 25.0 and 50.0 µg/mL of betacyanin fraction significantly downregulated NP at both the mRNA and protein levels. Betacyanin fraction of 25.0 µg/mL downregulated mRNA and protein NP by 1.078 (p-value = 0.0037) and 1.045-fold (p-value = 0.0275), respectively, while 50.0 µg/mL of betacyanin fraction downregulated mRNA and protein NP by 1.287 (p-value = 0.0018) and 1.519-fold (p-value = 0.0075), respectively. Interestingly, the positive control significantly downregulated protein NP expression level by a fold-change of 1.589 (p-value = 0.0062) compared to the untreated group but upregulated mRNA NP level by a fold-change of 1.112. Comparing between betacyanin-treated groups and the positive control, 25.0 (p-value = 0.0025) and 50.0 µg/mL (p-value = 0.0013) of betacyanin fraction demonstrated a significant difference only at the mRNA level and not at the protein level, as the positive control group showed upregulation at the mRNA level.

**Fig 4.**
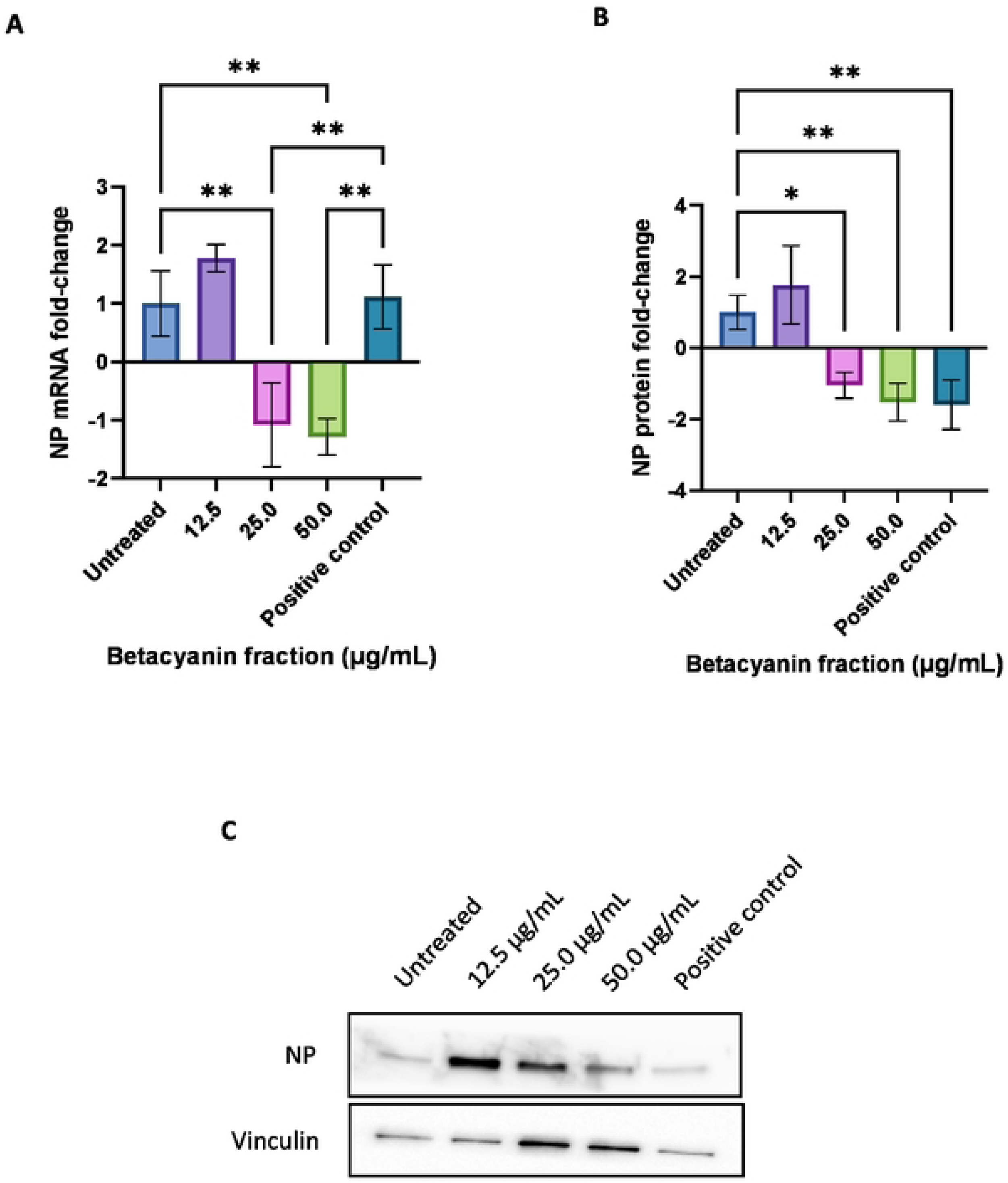
Fold changes in mRNA (A) and protein (B) NP expression levels and representative western blot image (C, supplementary information, Figure S10 & Sl l) in lA V-infected A549 cells after treatment with various betacyanin fraction concentrations. A549 cells were first seeded in a 12-well plate and then infected with an MOI of I with lAV. The cells received various concentrations of betacyanin fraction after infection for 24 h. The positive control group received I00 µM of oseltamivir phosphate. After 24 h, total RNA and protein were harvested from the cells using TRizol reagent. RNA was subjected to reverse transcription, qPCR, and data were analyzed using the delta-delta Ct method. Protein was subjected to SDS-PAGE and western blotting, and data was analyzed using densitometric analysis. Data were presented as means (n = 3) ± standard deviations and were subjected to one-way ANOVA to determine if changes in expression levels between groups were significant. Statistical significance was denoted by an asterisk (*), where * has a p-value < 0.05, and** has a p-value < 0.01.

IAV NP represents an important target as it is an essential factor in the virus life cycle, playing an important role in viral transcription and translation [28]. Multiple studies have shown that NP interacts and hijacks many host factors to favor its viral growth and survival [20, 21]. Preventing the virus from interacting with key host factors reduces virus titers *in vitro* and thus achieves antiviral effects [20, 21]. Additionally, NP has been shown to be a druggable target, and targeting NP has been shown to reduce viral titers *in vivo* [29]. Thus, measuring NP expression levels is a legitimate method to measure antiviral responses and effects. In this study, 25.0 and 50.0 µg/mL of the betacyanin fraction significantly downregulated NP at both mRNA and protein expression levels, as well as reducing virus titer preventing viral CPE formation.

### Betacyanin reduced virus titer and CPE formation after 48 h, with continued reduction in virus titer after 72 h in IAV-infected A549 cells

After treating infected cells with 50 µg/mL of betacyanin for 48 and 72 h, virus titers were found to be at 2587 and 203 pfu/mL, respectively (Fig 5a), or a reduction of 24% and 43% in virus titer, respectively when compared to the untreated group (Fig 5b). A significant reduction in virus titer was observed for both 48 and 72 h treatment groups (p-value = 0.0004 and 0.0334, respectively). Similar to the 24 h treatment, both time-points directly reduced virus titers in IAV-infected cells, achieving an antiviral effect. On the other hand, the virus titer of the positive control after 48 and 72 h was found to be 75 and 42 pfu/mL (Fig 5a), or a reduction of 98% and 88% in virus titer, respectively, when compared to the untreated group (Fig 5b). The number of plaques formed is visually decreased for both time-points (Supplementary information, Fig S12a, b, d, e).

**Fig 5.**
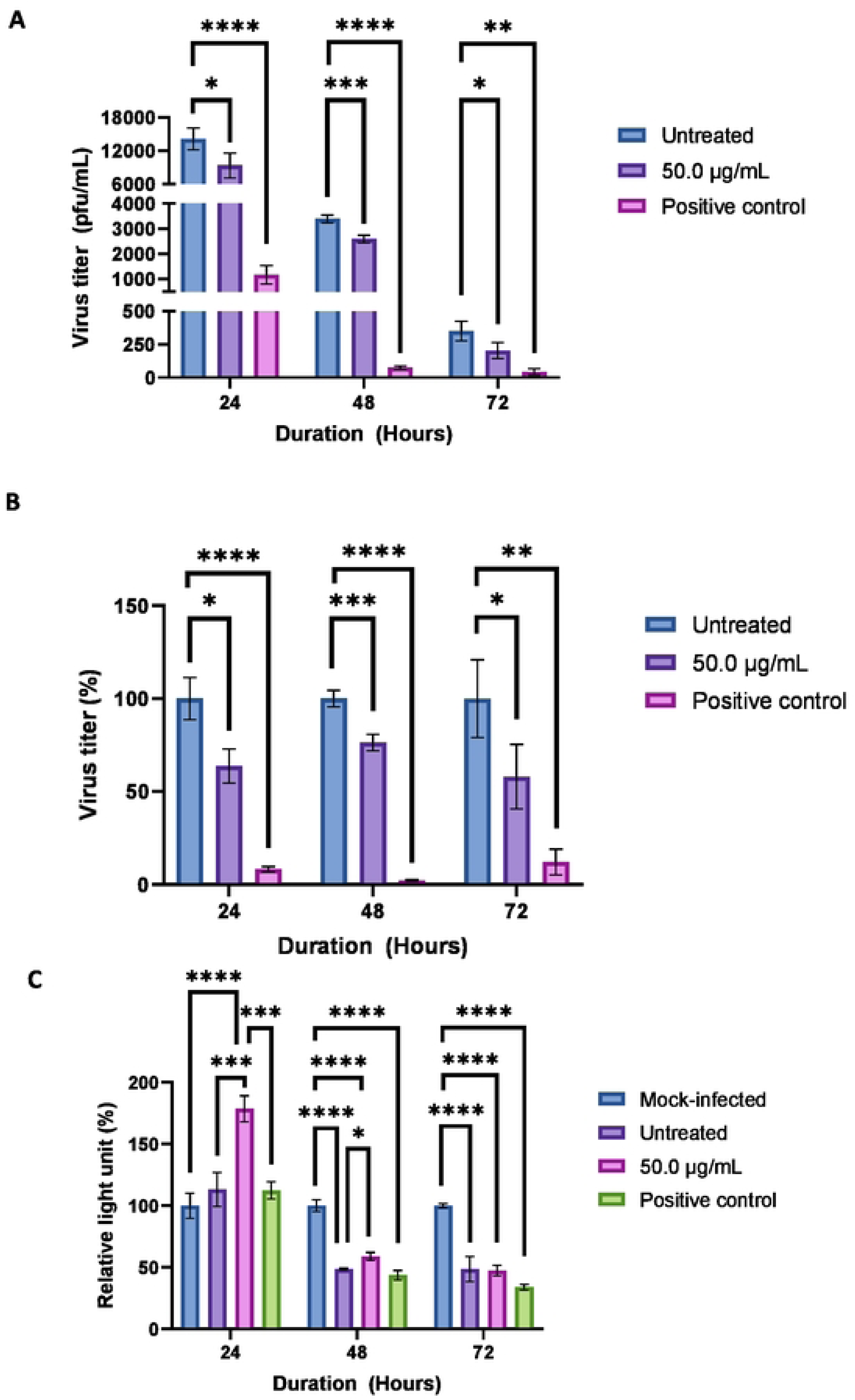
Virus titer in pfu/mL (A) and percentages **(B),** and relative light unit (RLU) after normalizing to the mock-infected group (C) in !AV-infected A549 cells after treatment with 50 µg/mL of betacyanin fraction for 24, 48, and 72 h. For Figures Sa and b, A549 cells were first seeded in a 12-well plate and then infected at an MO! of I with IAV. The cells received SO µg/mL ofbetacyanin after infection for 48 or 72 h. The positive control group received 100 µM of oseltamivir phosphate. The supernatant was harvested and subjected to plaque assay. For Figure Sc, A549 cells werefirst seeded in a 96-well plate and then infected at an MO!of I with IAV. The cells received 50µg/mL of betacyanin fraction after infection for 48 and 72 h. The positive control group received I00 µM of oseltamivir phosphate. All groups received virus infection, while the mock-infected group received virus growth med.ium only without the virus. After 48 or 72 h, the treatment media were removed and replaced with ATP Detection Reagent. RLU was acquired via a luminescence method. Data were presented as means (n = 3) ± standard deviations and were subjected to one-way ANOVA to determine if changes in virus titers or RLU between groups were significant. Statistical significance was denoted by an asterisk (*), where * has a p-value < 0.05, ** has a p-value <0.01, *** has a p-value of<0.001, and**** has a p-value of<0.0001.

Next, for the 48 h group, infected cells that received 50 µg/mL of betacyanin for 48 h observed a significant increase in RLU of 10.5% compared to the untreated group (p-value = 0.0270) (Fig 5c). An increase in RLU indicated reduced viral CPE formation. On the other hand, infected cells that received 50 µg/mL of betacyanin for 72 h experienced no significant changes in RLU (Fig 5c). This was a detrimental progression from the 24 h time-point, as a decrease in RLU can be observed. The positive control group that received oseltamivir phosphate for 48 and 72 h also did not experience significant changes in RLU compared to the untreated group. Looking from a disease progression perspective, comparing to the 24 h treatment time-point (Fig 5c), the change in RLU seemed to plateau after 48 h for the untreated group, as there was practically no change in RLU percentages between the 48 and 72 h (48.5% and 48.6% respectively), indicating no further CPE formation (Fig 5c).

Morphologically, cell debris and cellular damage can be observed for the untreated group after both 48 and 72 h (Supplementary information, Fig S13 & S14). Interestingly, cell debris was not observed for infected cells that received 50 µg/mL of betacyanin for 48 and 72 h, but morphological changes in the form of CPE formation can be observed (Supplementary information, Fig S13 & S14). This was a detrimental progression from the 24 h treatment, as previous results showed that 12.5, 25.0, and 50.0 µg/mL of betacyanin significantly prevented CPE formation in both quantitative and qualitative measures (Fig 2 & S8).

Putting both plaque assay and CPE formation results together, 50 µg/mL betacyanin was able to consistently reduce virus titers from infected cells to up 72 h, however, CPE formation was only reduced up to48 h, and no changes in CPE formation were observed after 72 h. These results indicate that future virus production from infected cells will be reduced until 72 h. However, cellular damage in the form of CPE can only be reduced until 48 h. This study demonstrated that 12.5, 25.0, and 50.0 µg/mL of betacyanin fraction demonstrated antiviral activity against IAV-infected A549 cells by preventing viral CPE formation and lowering virus titer, with the 50 µg/mL of betacyanin fraction exhibiting antiviral activity up to 72 h. Furthermore, 25.0 and 50.0 µg/mL of betacyanin downregulated a key IAV factor, NP, at both mRNA and protein levels. While the positive control was better at reducing virus titer, betacyanins seemed to possess other protective effects by preventing viral CPE formation that assisted in cell survival, suggesting further putative interactions with other host factors in IAV-infected A549 cells, thereby opening new avenues for studies on host-viral relations with betacyanins.

## Supporting information

S1 – 7 Fig. LC-MS and LC-MS/MS spectra of phyllocactin, betanin, isobetanin, and hylocerenin.

S8 Fig. CPE microscopic images, 24 h.

S9 Fig. Representative plaque images, 24 h.

S10 Fig. Representative western blot image of NP, 24 h.

S11 Fig. Representative western blot image of vinculin, 24 h.

S12 Fig. Representative plaque images, 48 and 72 h.

S13 Fig. CPE microscopic images, 48 h.

S14 Fig. CPE microscopic images, 72 h.

## Author contributions

Conceptualization: Wee Sim Choo

Data curation: Chie Min Lim

Formal analysis: Chie Min Lim

Funding acquisition: Wee Sim Choo

Investigation: Chie Min Lim

Project administration: Wee Sim Choo

Resources: Wee Sim Choo

Supervision: Wee Sim Choo, Sunil Kumar Lal

Writing – original draft: Chie Min Lim

Writing – review & editing: Wee Sim Choo, Sunil Kumar Lal, Nurulfiza Mat Isa, Abdul Rahman Omar

## Data availability

All data are fully available without restriction.

